# GANscan: continuous scanning microscopy using deep learning deblurring

**DOI:** 10.1101/2022.02.22.481502

**Authors:** Michael John Fanous, Gabriel Popescu

**Affiliations:** Quantitative Light Imaging Laboratory, Beckman Institute for Advanced Science and Technology, University of Illinois at Urbana-Champaign, Urbana, Illinois 61801, USA; Department of Bioengineering Department of Bioengineering, University of Illinois at Urbana-Champaign, 306 N. Wright Street, Urbana, IL 61801, USA; Department of Electrical and Computer Engineering, University of Illinois at Urbana-Champaign, 306 N. Wright Street, Urbana, IL 61801, USA

## Abstract

Most whole slide imaging (WSI) systems today rely on the “stop-and-stare” approach, where, at each field of view, the scanning stage is brought to a complete stop before the camera snaps a picture. This procedure ensures that each image is free of motion blur, which comes at the expense of long acquisition times. In order to speed up the acquisition process, especially for large scanning areas, such as pathology slides, we developed an acquisition method in which the data is acquired continuously while the stage is moving at high speeds. Using generative adversarial networks (GANs), we demonstrate this ultra-fast imaging approach, referred to as GANscan, which restores sharp images from motion blurred videos. GANscan allows us to complete image acquisitions at 30x the throughput of stop-and-stare systems. This method is implemented on a Zeiss Axio Observer Z1 microscope in brightfield mode, requires no specialized hardware, and accomplishes successful reconstructions at stage speeds of up to 5,000 μm/s. We validate the proposed method by imaging H&E stained tissue sections. Our method not only retrieves crisp images from fast, continuous scans, but also corrects any defocusing that occurs during scanning. Using a consumer GPU, the inference runs at <20ms/ image.

## Introduction

Numerous microscopy applications require large fields of view (FOV), including digital pathology ^1^, micro-mirror and biosensor assembly ^2^, and *in vivo* imaging ^3^. Acquisition time demands are a major bottleneck to fixing modest or partially filled FOVs in standard microscopy techniques. Improvements in both hardware and computation are thus actively sought to push the efficiency of optical measurements beyond traditional boundaries. Accelerating either image acquisition or analysis can have drastic benefits in diagnostic assessments and has been shown to provide critical advantages in cell detection^4^, disease screening^5^, clinical studies^6^ and histopathology^7,8^.

In standard microscope systems, the objective lens dictates the resolution and field-of-view (FOV), forcing a trade-off between the two parameters. In commercial whole slide scanners, the FOV is extended through lateral scanning and image mosaicking. Most forms of microscopy require serial scanning of the sample region, which slows down measurement acquisitions and diminishes the temporal resolution.

There are three classes of strategies used in traditional microscopy for slide-scanning. The first technique uses the so called “stop-and-stare” style, which entails sequentially moving the sample across a scanning grid, pausing the stage, and exposing the camera for discrete recordings. This tactic generates high-quality images as a result of long measurement durations, but is not especially time-efficient^9^. A second technique involves illuminating a moving sample with bursts of light that help circumvent the motion blur, which would otherwise compromise the image resolution. As a result of the short exposure times with this method, the resulting images have a relatively poor signal-to-noise ratio (SNR)^9^. Thus, there is a cost to optimizing image clarity or acquisition speed in these approaches. Third, there are line scanning ^10^ and time-delay integration (TDI) ^11^ methods, which use 1D sensors, where a camera vertically handles continuous signals line by line to reduce read-out time and increase SNR. However, even the latest versions of these instruments require specialized imaging equipment and readout methods^12,13^.

Different imaging methods have been proposed to improve the throughput of scanning-based microscopy techniques, such as multifocal imaging^14^ and coded illumination^9^. Computational methods of microscopy imaging^15–19^, such as ptychography, which scans and fuses portions of spatial frequencies, can produce large FOVs with resolutions that surpass the objective’s diffraction limit. However, these solutions end up either complicating the microscopy system configuration, deteriorating the image quality, or extending the post-processing period. Additionally, iterative algorithms that are used in Fourier ptychography to reconstruct an image from a sequence of diffraction patterns often suffer from convergence issues^20^.

The mechanical specifications of the scanning stage, rather than the optical parameters of the microscope, generally hinder the throughput performance of WSI systems^21^. The space-bandwidth product (SBP), which is the dimensionless product of the spatial coverage (FOV) and the Fourier coverage (resolution) of a system, can quantify the information across an imaging system^22^. Enhancements to the SBP have been the objective of various innovations in imaging techniques ^23–28^, but typically require either specialized hardware or time-consuming post-processing.

The advent of accessible deep learning tools in recent years has led to a new host of strategies to address lingering microscopy challenges ^27^, including super-resolution imaging ^29^, digital labeling of specimens^30–39^, Fourier ptychography microscopy^26^, and single-shot autofocusing^40^, among others^41^. These methods, which take advantage of recent breakthroughs in deep learning, need no modification to the underlying microscopic gear and produce faster and more comprehensive imaging results than traditional image reconstruction and post-processing algorithms. Generative adversarial networks (GANs), which comprise two opposing networks competing in a zero-sum dynamic, have been especially prominent in image-to-image translation tasks, due in large part to their outstanding execution of pixel-to-pixel conversions^31,42^.

In this work, we propose a computational imaging technique, termed GANscan, which employs a GAN model to restore the spatial resolution of blurred videos acquired via continuous stage scanning at high speeds using a conventional microscopy system. Our method involves continuously moving the sample at a stage speed of 5,000 μm/s and an acquisition rate of 30 frames per second (fps). This acquisition speed is on par with the state-of-the-art TDI technology of 1.7-1.9 gigapixels in 100 seconds ^11,13^. However, unlike TDI, our approach is using standard optical instrumentation, which lowers the threshold for broad adoption in the field.

In contrast to other high-throughput imaging endeavors, GANscan adds no complexity to the hardware, with single frame restorations that can be computed in a matter of milliseconds. The results of this novel technique demonstrate that basic modifications in measurements, coupled with artificial intelligence (AI), can provide the framework for any rapid, high-throughput scanning operation.

This paper is structured as follows: first, we present the workflow for continuous imaging microscopy in both slow and fast acquisitions. Second, we describe the theory behind blur motion artifacts and why deconvolutions are limited in restoring the spatial bandwidth of control images. Third, we discuss the imaging procedures and registration of slow-moving samples with the motion-smeared ones. Fourth, the parameters of the GANscan network are explained, as well as the data processing techniques prior to model training. Lastly, reconstruction performances are evaluated using an unseen test set, which is also compared against deconvolutions using standard image metrics.

## Results

### Workflow

Figure 1 depicts the workflow of our approach. To demonstrate the benefits of this technique, we imaged a large sample of a pathological slide of a ductal carcinoma in situ (DCIS) biopsy, covering roughly half a standard microscopy slide area (∼30 mm x 15 mm). The slide was scanned in a row-major configuration, capturing movies across the slide horizontally (Fig. 1a). There were no modifications to the standard commercial brightfield microscope (Axio Observer Z1, Zeiss), and the only adjustments in the measurement were the speed of the stage and the continuous recording of the camera. In order to obtain ground truth images for training, the same rows were captured at a slow (50 μm/s) stage speed and at the same exposure time of 2 ms. Once pairs of sharp and defocused images were assembled through Pearson correlations, a GAN network was trained to enable restoring unseen motion blurred micrographs (Fig. 1b).

**Figure 1.**
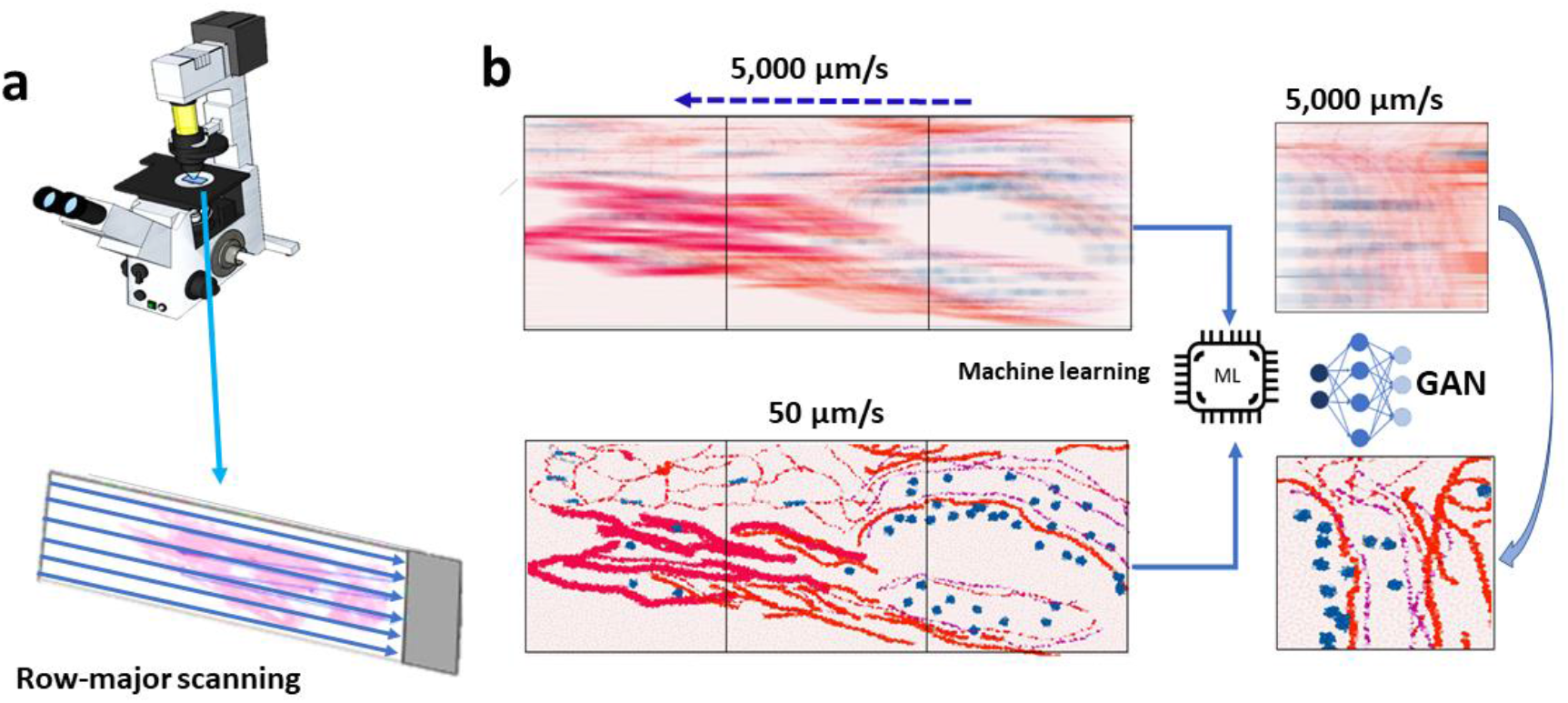
**a** Scanning stage of the AXIO observer Zeiss microscope with an example slide showing the row-major continuous scanning direction. **b** Motion blurred reconstruction scheme using a slow-moving stage as the control for GAN training.

### Theory

At rest, let the image be *I*(*x, y*). During the sample translation, the translated image, *<u>I</u>*, has the following time dependence:

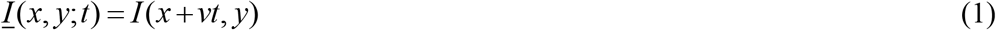

where *v* is the stage speed. Considering the camera integration time T, the “blurred” detected frame is then:

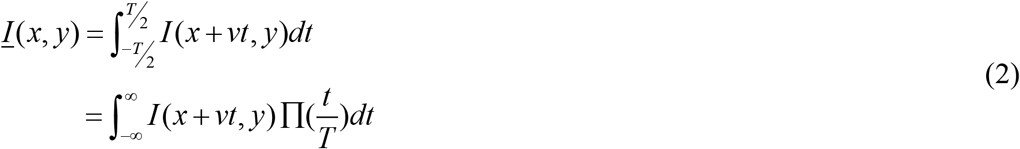

where 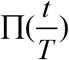 is the 1D rectangular function of width *T*. The integration is the sum of the frames accumulated during the acquisition time *T* (Fig. 2a).

**Figure 2.**
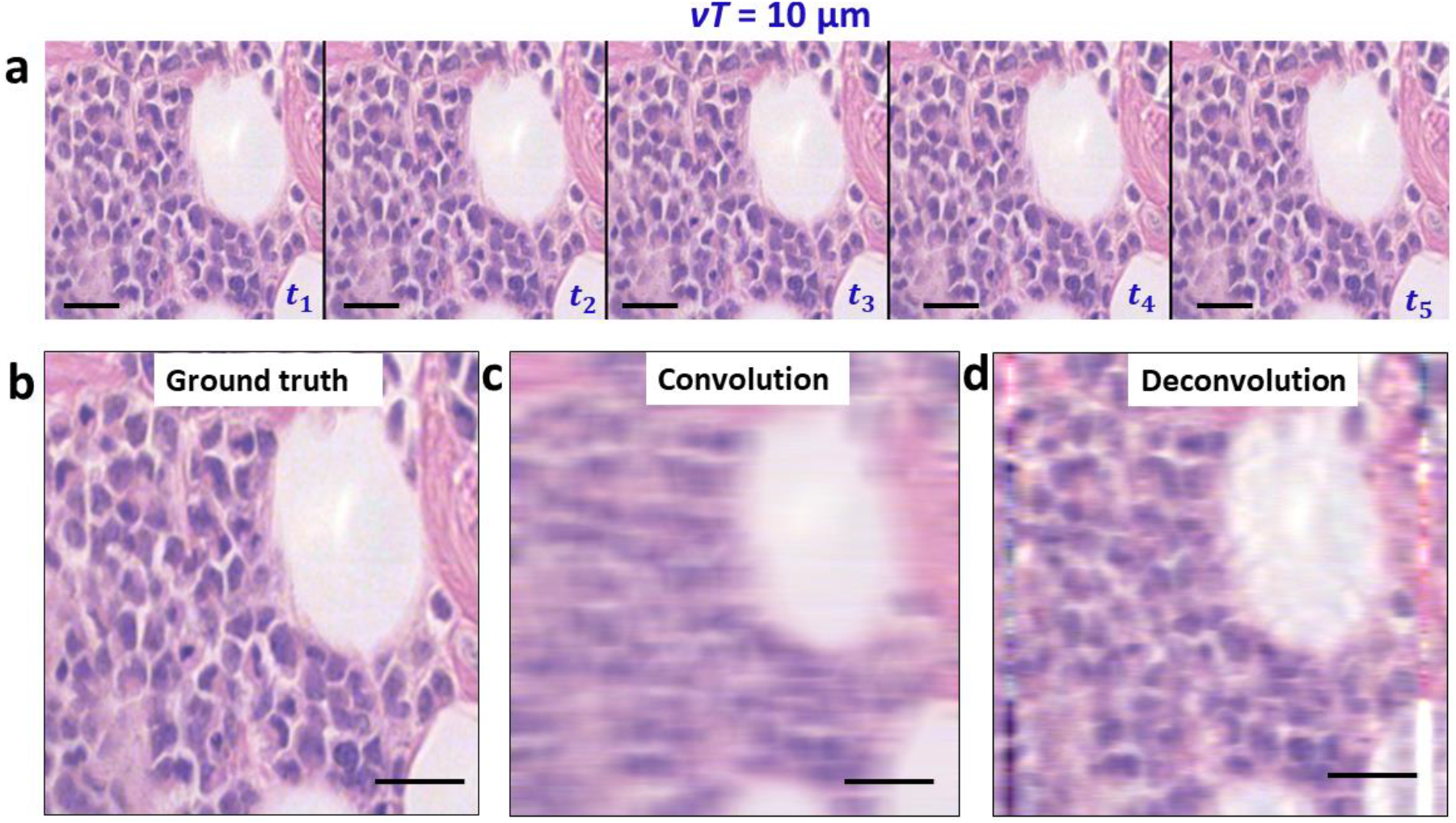
**a** A few frames from a video taken with a stage speed of 50 μm/s, with time labels indicating a slow forward movement. **b** Middle image of the sharp sequence **a** used as ground truth. **c** The convolution of **b** with the blur function. **d** The deconvolution of **c** with the blur function. Scale bar **25 µm**.

Using the central-ordinate theorem ^43^:

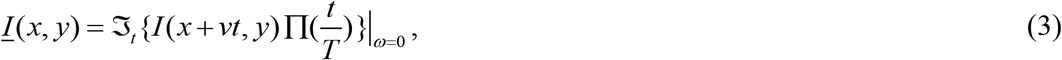

where ω is the angular frequency. Since 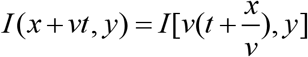, the temporal Fourier transform reads

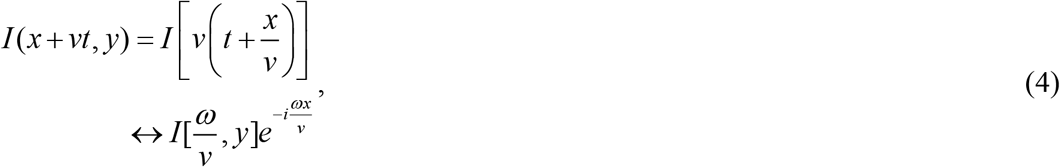

where ↔ indicates the Fourier transformation.

Using the convolution theorem ^44^, Eq. 3 can be rewritten as:

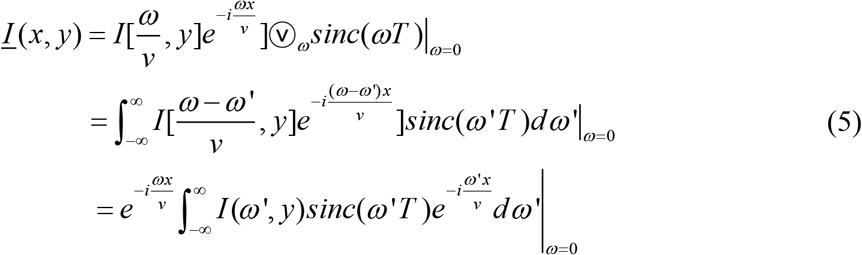

In Eq. 5, we recognize a Fourier transform of a product, which yields the following convolution operation,

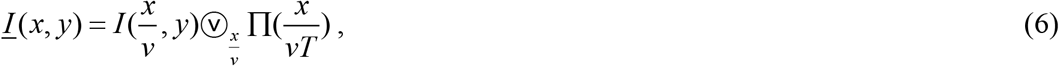

where 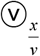 indicates the convolution operator over the variable *x/v*, which has dimensions of time. This result captures the physical description of the image *spatial* blurring as the result of a *temporal* convolution operation. Thus, the smeared image is the sharp image convolved along the direction of the scan by a rectangular function, which has a width proportional to the acquisition time. For a scanning speed of *v* = 5, 000 *μm*/*s* and *T* = 2 *ms, vT* = 10 *μm*. This corresponds to a length roughly twenty times the diffraction resolution of our imaging system.

### Deconvolution

We performed the 1D *deconvolution* on our acquired images, thus, inverting the effect of Eq. 5, and used the results as the standard of comparison for the deep learning results. These deconvolutions were evaluated by first establishing the best match through the ‘convolve’ filter in ImageJ, and then using the same line dimension in MATLAB with the ‘deconvblind’ function. This tool deconvolves an image via the maximum likelihood algorithm and a starting estimate of the point-spread function (PSF), which in our case is a single row of 47 pixels of value 1.

A sample frame of the biopsy and its convolution with the line of the blur width are shown in Figs.2 b-c, and deconvolving again produces the original frame but with compromised high spatial frequencies (Fig.2d). The artifacts of lines along both edges of the image are a result of the filter brushing against the boundaries of the image. The deconvolution operation succeeds at shrinking features horizontally to restore their true width. However, the image still suffers from poor overall resolution, due in part to the higher spatial frequencies being permanently lost through the convolving effect of imaging a rapidly moving sample. This shortcoming is our principal motivation of employing deep learning techniques to predict the standard spatial bandwidth.

### Image pair registration

In order to prepare pairs of blurred and sharp images for training, consecutive sharp images in the fast videos were matched to their motion-smeared counterparts by evaluating the maximum Pearson correlations in a set of slightly shifted clear images (Figs. 3, S1). The “ground truth” images were captured at a stage speed of 50 μm/s, which, at the acquisition time of 2 ms results in a blur size of 0.1 *μ*m, *i*.*e*., below the diffraction limit of our system. As a result, there are approximately 100 frames in the sharp videos for each image in the 5,000 μm/s, motioned blurred videos, as shown in Figure 3a.

**Figure 3.**
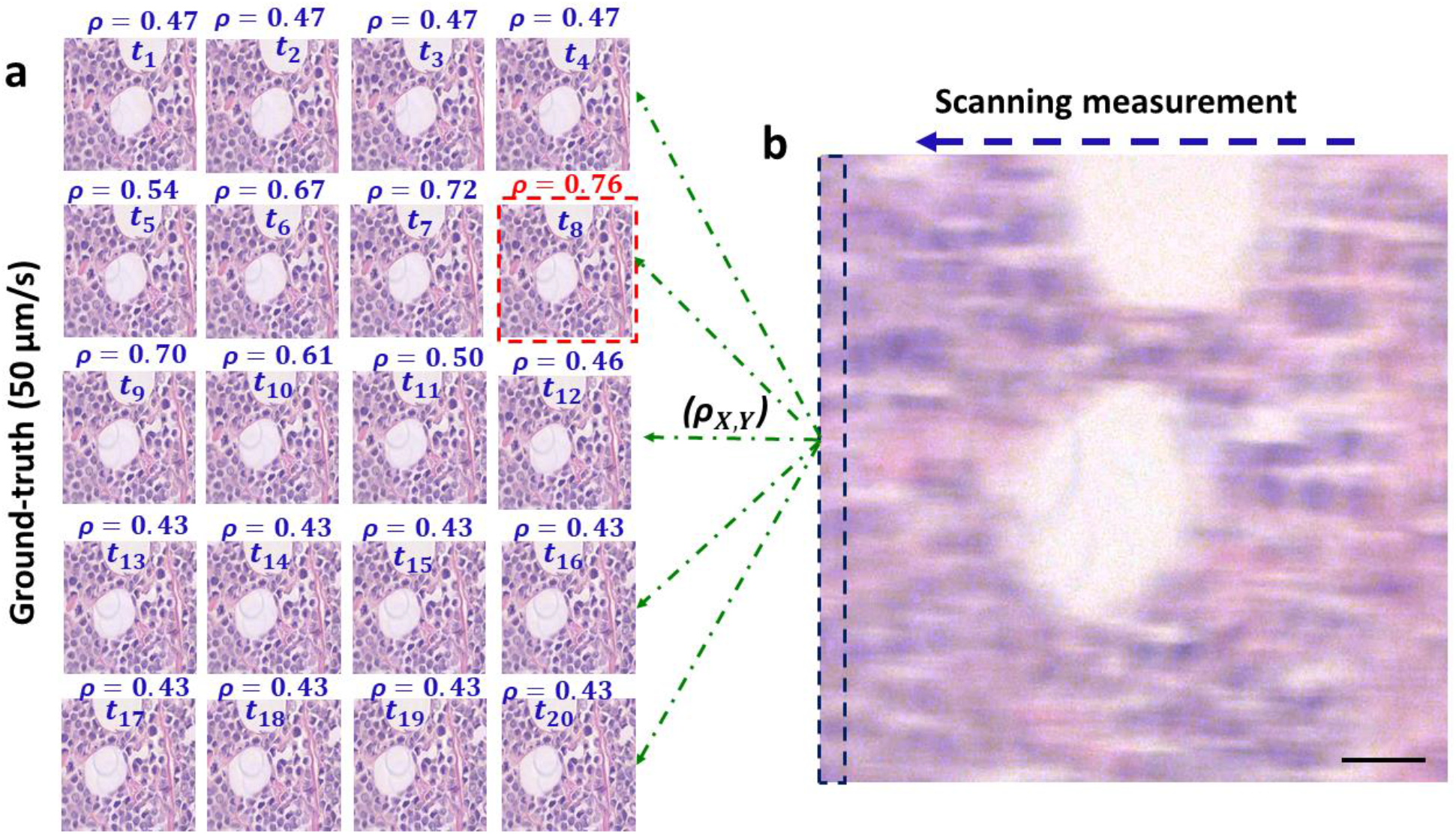
**a** Registration of images through maximum Pearson coefficient between sharp frames at 50 μm/s and **b** the blurry one at 10,000 μm/s. The scanning measurement is compared with each sharp contender (green dashed lines) and the resulting coefficient score is listed above each image. The best match is delineated in red, indicating the sharp image bearing the overall highest resemblance to the streaked measurement. The dashed rectangle at left edge of **b** outlines the extra margin of features contained in a blurry image (10 μm).

Evaluating the Pearson correlation between the input (smudged) frame and a series of potential ground truth frames produces values ranging from 0.4 to 0.76. The frame associated with the highest Pearson correlation value was selected as the ground truth. It should be noted that the rapidly captured images expose the camera to a larger field of view than the slowly-captured ones by the length of the blur, which is 10 *μ*m for a scanning speed of 5,000 μm/s. This difference is delineated in Figure 3b.

### Generative adversarial network (GAN)

Once the registered pairs were assembled, they were cropped and resized to dimensions of 256×256×3 (3 RGB color channels) for faster computation, with 1,050 images earmarked for training and 50 reserved for testing. The architecture of the model consists of a generator U-net with eight encoding and decoding layers, and a four downchannel discriminator, all displayed in Figure S3. As shown in Figure 4, the network input is the motion-blurred image and the control is the slowly scanned, sharp image. Since the slide was scanned in a row-major style, the margin of additional field of view is always on the same side, which is likely to help the network to undo the motion distortion. The GAN model was trained for over 200 epochs (Fig. 4c) until the loss function plateaued. Our results indicate that running the model on the training set produces nearly perfect restorations (Fig. 4d). The spatial power spectrum of the input image (Fig 4e) clearly shows a smaller range of higher spatial frequencies than that of the restored image (Fig 4f). Interestingly, the power spectrum of the input image has higher spatial frequencies along the vertical axis as a result of the smearing produced along the x-axis, whereas the power spectrum of the restored image is broader and more isotropic.

**Figure 4.**
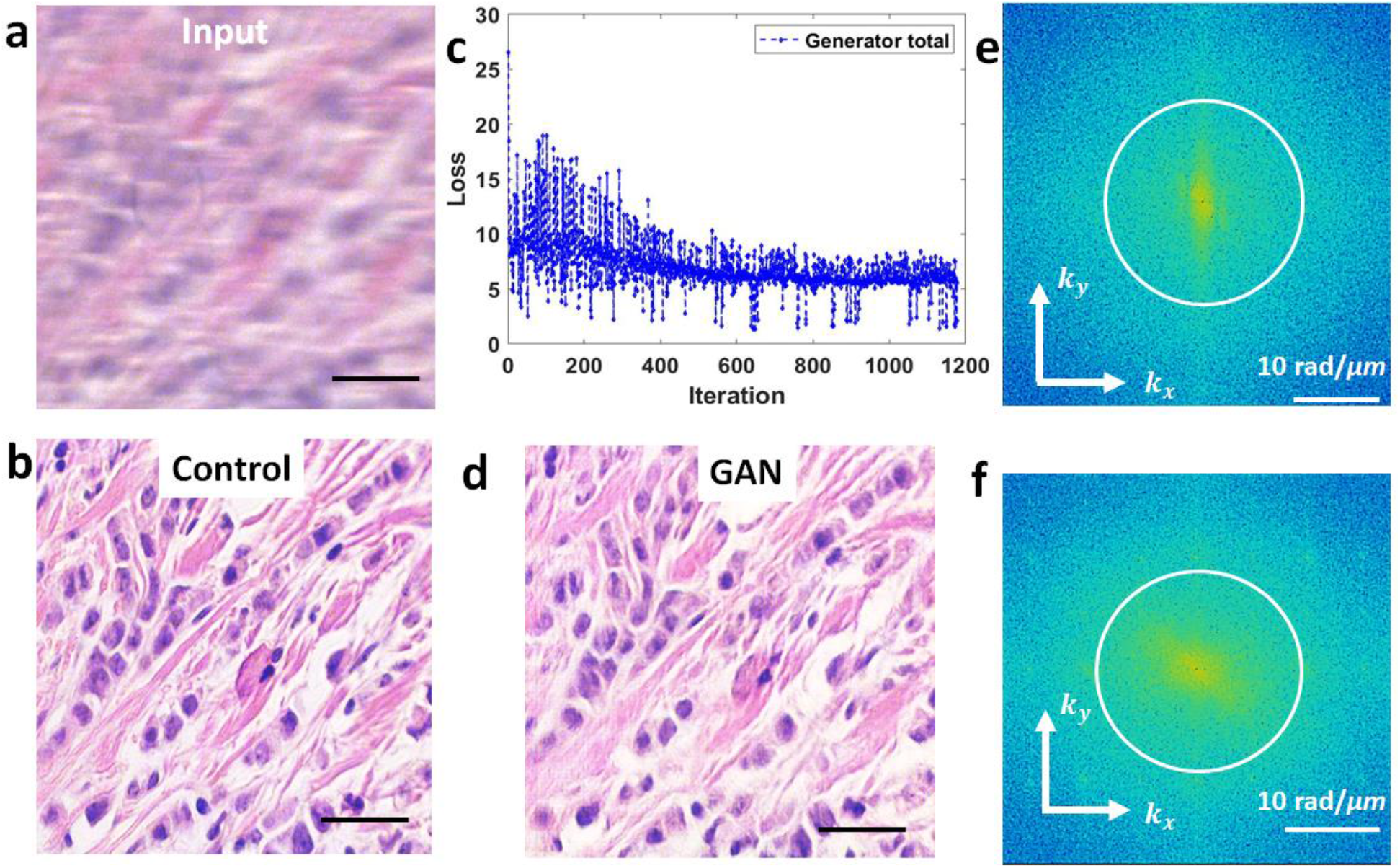
**a** Training example of the blurry input, **b** and control, **c** with the loss plot. **d** The generated result for a training instance. **e** The power spectra of the input and **f** of the result with a circle outlining the diffraction spot. Scale bar in **a, b**, and **d 25 µm**.

### Performance testing

Once the training was complete, the model was tested on 50 *unseen* images. A sample of these are displayed in Figure 5. The network does an effective job at restoring the high spatial frequencies of epithelial and stromal (fibrous) areas, as compared to the line deconvolutions (Fig. S2a). Since the cellular and fibrous areas are recovered with such high fidelity, the diagnostic information in the tissue images is maintained in full. In terms of numerical assessments, the test sets achieved an average structural similarity index measure (SSIM) of 0.82 and a mean peak signal-to-noise-ratio (PSNR) of 27 when calculated against their controls. For the same dataset, the deconvolution results gave inferior results of SSIM and PSRN of 0.71 and 26, respectively.

**Figure 5.**
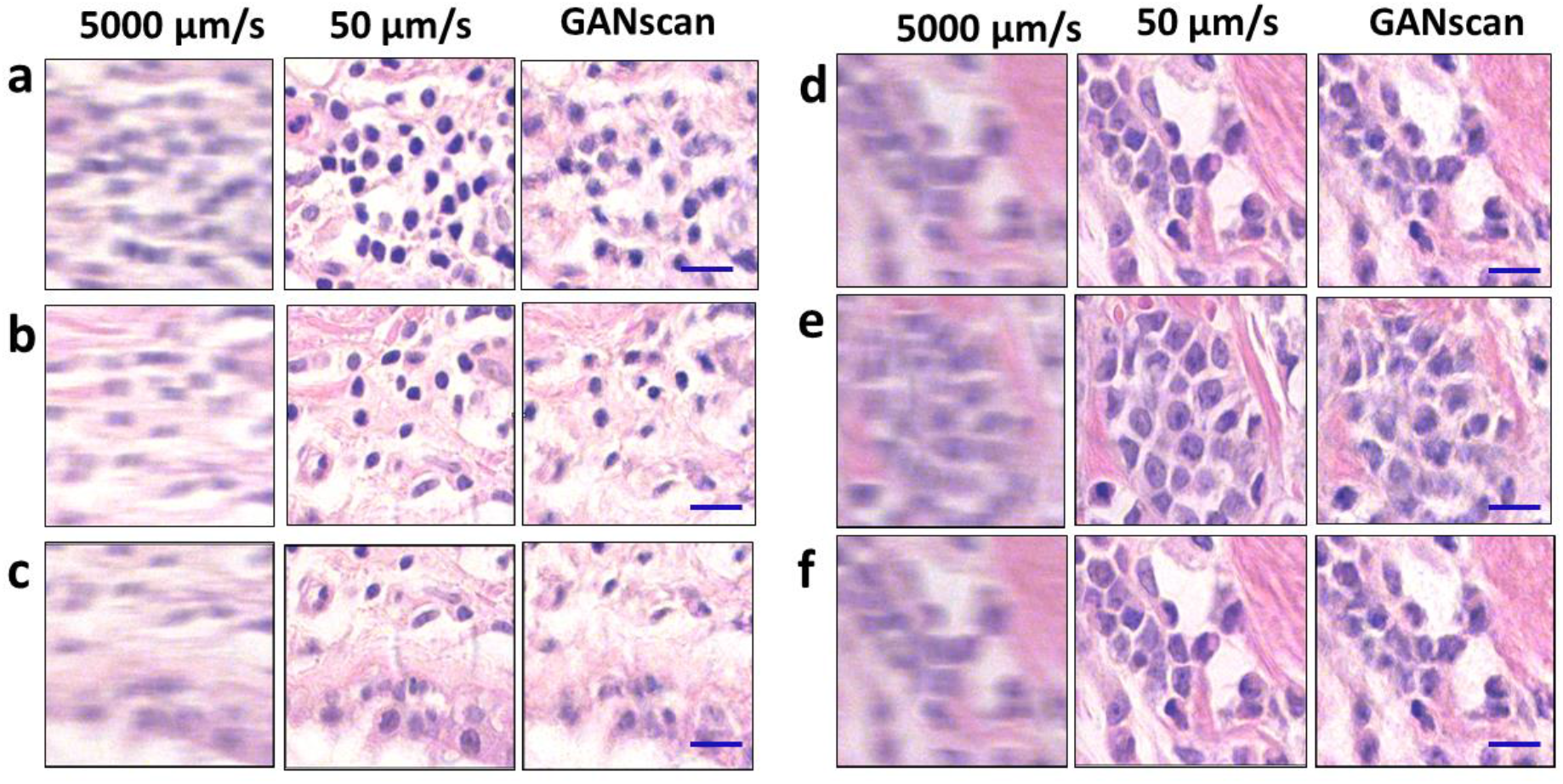
**a-f** Test set conversion of 5000 µm/s with control of 50 µm/s. Scale bar **25 µm**.

Large mosaics of the motioned blurred images were also reconstructed (Fig. 6) by concatenating the images horizontally and vertically in their respective scanning order, producing a 7 × 15 stitch of roughly 3 mm x 1.5 mm in size. The difference in clarity is much less apparent with such a large FOV, but at a closer look it is evident there is significant improvement in the overall distinction of features. Stitches for 4,000 μm/s were also made for comparison (Fig. S4).

**Figure 6.**
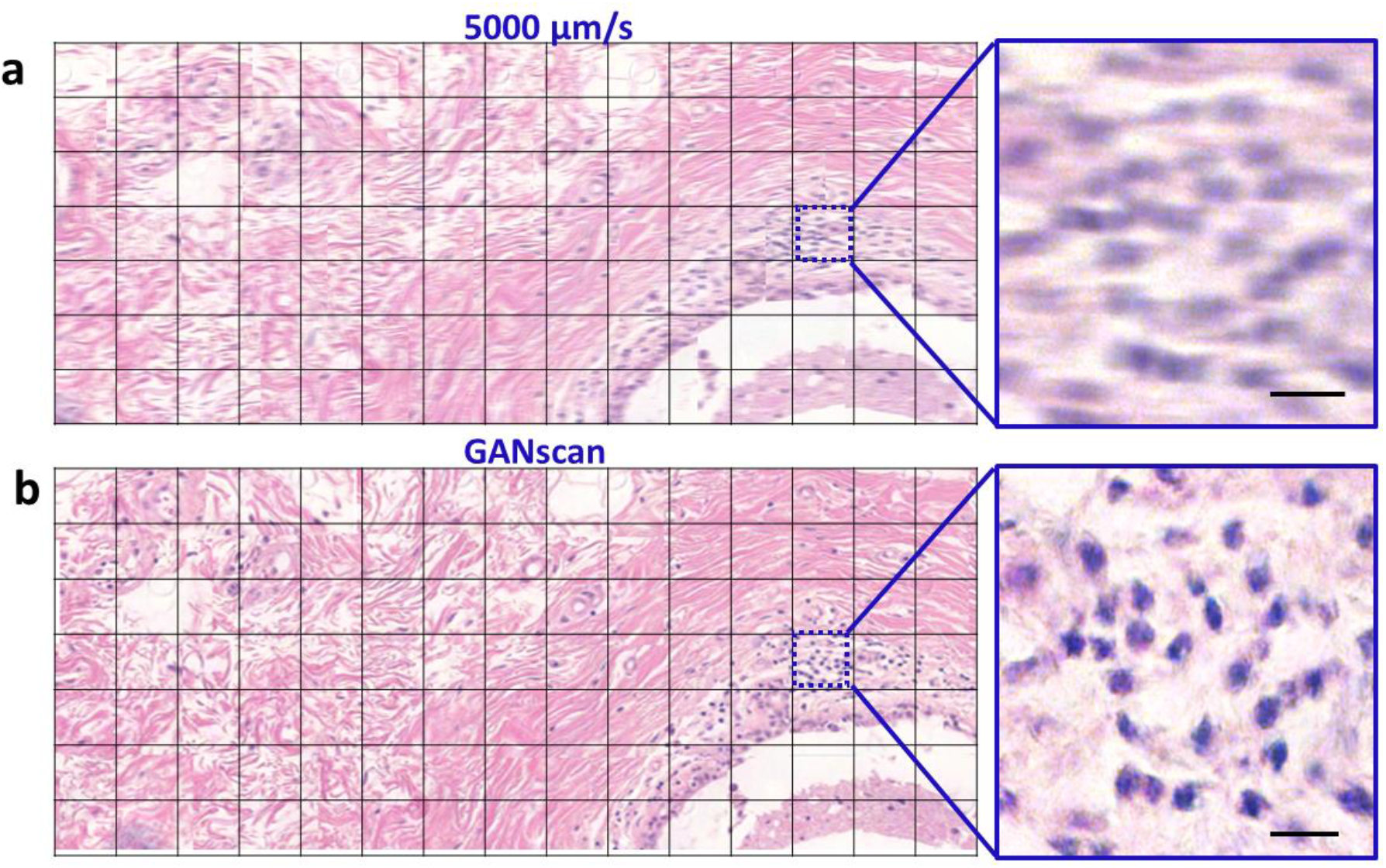
**a** Stitch of the motion blurred images, **b** as well as the reconstructed GANscan, showing a large area of a breast biopsy, with respective zoom-ins. Scale bar **25 µm**.

## Discussion

We presented a high-throughput imaging approach, GANscan, which employs continuous motion deblurring using labelled GAN reconstruction. Through both theoretical and experimental analysis, we have demonstrated the applicability of our method to brightfield microscopy on tissue slides. Our results indicate that GAN models provide, in combination with greater stage speeds, up to 30x faster acquisition rates than in conventional microscopy. This throughput is superior or on par with the state-of-the-art rapid scanning techniques, which in turn use nonstandard hardware. GANscan requires no specialized equipment and generates restored images with successfully removed motion blur. Of course, should a camera with a higher frame rate be used, the stage speed can be scaled up proportionally. Further, our proposed deep learning deblurring method produces high-quality reconstructions which restores the high frequency portions of the tissue, as opposed to deconvolution operations.

Such a methodology will not only provide a drastic benefit in the clinical setting for pathologists, but at the research level as well, including cell cultures of large dimensions. Future work should address achieving similar results with different microscope modalities, such as fluorescence and quantitative phase imaging.

## Material and methods

### Image acquisition

Images were acquired with a commercial brightfield microscope (Axio Observer Z1, Zeiss) and a Point grey color camera, using a Zeiss EC Plan-Neofluar 40x/0.45 NA objective. The sample is a ductal carcinoma in situ (DCIS) breast tissue biopsy. The stage speed and coordinates were precisely manipulated using the Zeiss MTB (MicroToolBox) software, and the camera settings, such as shutter time (2 ms), frame rate (30 pfs), and gain (8 dB), were selected using the Grasshopper GRAS-2054C software. For stitching images, a vertical step size of 200 µm was used, and horizontal videos were acquired for 1 minute at the slow speed of 50 µm/s to ensure the correspondence of 15 horizontally adjacent frames in the video captured at 5000 µm/s. The video of each row at the accelerated stage speed was 0.6 seconds. After the image acquisition, off-line processing involved image registration of blurry and sharp images through MATLAB with Pearson correlation estimates. For the 5,000 µm/s datasets, we extracted 256×256 crops from paired images to create a training volume of 1050 image pairs.

We performed deconvolutions on each input test image and compared them with GANscan results, as shown in Figure S1. The mean SSIM of the GANscan images is 0.82, while the deconvolved images had an SSIM of 0.73, when compared to the same control images. PSNR values were also calculated with GANscan outperforming deconvolutions 27 to 26. All analysis was performed in MATLAB.

### Machine learning

The conversion of motion blurred micrographs to sharp images was accomplished using the conditional generative adversarial network (GAN) pix2pix (Fig. S3) ^45^. 1050 blurry and sharp brightfield RGB image pairs were passed through the network. Original dimensions of the micrographs were 600×800×3 pixels. These were cropped and resized to 256×256×3 pixels before being trained on. The learning rate of the generator’s optimizer was 0.0002 and the minibatch size was set to 1. In this network, a generator (G) is trained to produce outputs that cannot be distinguished from ground truth images by a trained adversarial discriminator, *D*, which is designed to perform as well as possible at detecting the generator’s incorrect data ^45^. The GAN loss is one where *G* works to minimize the value while an adversarial *D* attempts to maximize it:

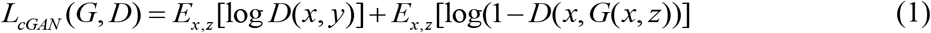

Where *E*_*x,z*_ is the anticipated value of all real and fake instances, x is the image, and z is the generated random noise. An L1 loss is then combined with this to generate the discriminator’s total loss function.

In order to confirm the accuracy of the translated images, we tested the model on 50 unseen images. Training was performed over 200 epochs, with datasets that were augmented beforehand through rotations and mirroring. Overall, the training took 7 hours, and the inference required less than 20 ms per image (256×256 pixels).

## Acknowledgements

The authors are grateful to Betsy Barnick and Priya Dutta at Carle Foundation Hospital for providing the pathology slides. This work was funded by the National Institute of Health (R01CA238191, R01GM129709).

## Data availability

All data required to reproduce the results can be obtained from the corresponding author upon a reasonable request.

## Code availability

All the code required to reproduce the results can be obtained from the corresponding author upon a reasonable request.

## Conflict of interest

There are no conflicts of interest.

## Notes

### Competing Interest Statement

The authors have declared no competing interest.

## References

1 Horstmeyer, R., Ou, X., Zheng, G., Willems, P. & Yang, C. Digital pathology with Fourier ptychography. Computerized Medical Imaging and Graphics 42, 38–43 (2015).

2 Potsaid, B., Bellouard, Y. & Wen, J. T. Adaptive Scanning Optical Microscope (ASOM): A multidisciplinary optical microscope design for large field of view and high resolution imaging. Optics express 13, 6504–6518 (2005).

3 Webb, R. H. & Rogomentich, F. Confocal microscope with large field and working distance. Applied optics 38, 4870–4875 (1999).

4 Alegro, M. et al. Automating cell detection and classification in human brain fluorescent microscopy images using dictionary learning and sparse coding. Journal of neuroscience methods 282, 20–33 (2017).

5 Brodin, P. & Christophe, T. High-content screening in infectious diseases. Current opinion in chemical biology 15, 534–539 (2011).

6 Messner, C. B. et al. Ultra-fast proteomics with Scanning SWATH. Nature biotechnology, 1–9 (2021).

7 Remmelinck, M. et al. How could static telepathology improve diagnosis in neuropathology? Analytical Cellular Pathology 21, 177–182 (2000).

8 Gareau, D. S. et al. Confocal mosaicing microscopy in Mohs skin excisions: feasibility of rapid surgical pathology. Journal of biomedical optics 13, 054001 (2008).

9 Phillips, Z. F., Dean, S., Recht, B. & Waller, L. High-throughput fluorescence microscopy using multi-frame motion deblurring. Biomedical optics express 11, 281–300 (2020).

10 Ho, J. et al. Use of whole slide imaging in surgical pathology quality assurance: design and pilot validation studies. Human pathology 37, 322–331 (2006).

11 Hamamatsu. High throughput imaging in low light applications. TDI Solutions (2011).

12 De Moor, P. et al. in 2014 IEEE International Electron Devices Meeting. 4.6. 1-4.6. 4 (IEEE).

13 Iftimia, N. V. et al. Adaptive ranging for optical coherence tomography. Opt. Express 12, 4025–4034 (2004).

14 Prabhat, P., Ram, S., Ward, E. S. & Ober, R. J. Simultaneous imaging of different focal planes in fluorescence microscopy for the study of cellular dynamics in three dimensions. IEEE transactions on nanobioscience 3, 237–242 (2004).

15 Abrahamsson, S. et al. Fast multicolor 3D imaging using aberration-corrected multifocus microscopy. Nature methods 10, 60–63 (2013).

16 Bouchard, M. B. et al. Swept confocally-aligned planar excitation (SCAPE) microscopy for high-speed volumetric imaging of behaving organisms. Nature photonics 9, 113–119 (2015).

17 Nakano, A. Spinning-disk confocal microscopy—a cutting-edge tool for imaging of membrane traffic. Cell structure and function 27, 349–355 (2002).

18 Li, H. et al. Fast, volumetric live-cell imaging using high-resolution light-field microscopy. Biomedical Optics Express 10, 29–49 (2019).

19 Martínez-Corral, M. & Javidi, B. Fundamentals of 3D imaging and displays: a tutorial on integral imaging, light-field, and plenoptic systems. Advances in Optics and Photonics 10, 512–566 (2018).

20 Hu, C., Kandel, M. E., Lee, Y. J. & Popescu, G. Synthetic aperture interference light (SAIL) microscopy for high-throughput label-free imaging. Applied Physics Letters 119, 233701 (2021).

21 Farahani, N., Parwani, A. V. & Pantanowitz, L. Whole slide imaging in pathology: advantages, limitations, and emerging perspectives. Pathology and Laboratory Medicine International 7, 23–33 (2015).

22 Lohmann, A. W., Dorsch, R. G., Mendlovic, D., Zalevsky, Z. & Ferreira, C. Space–bandwidth product of optical signals and systems. JOSA A 13, 470–473 (1996).

23 Gustafsson, M. G. Surpassing the lateral resolution limit by a factor of two using structured illumination microscopy. Journal of microscopy 198, 82–87 (2000).

24 Rodenburg, J. M. & Faulkner, H. M. A phase retrieval algorithm for shifting illumination. Applied physics letters 85, 4795–4797 (2004).

25 Tian, L., Li, X., Ramchandran, K. & Waller, L. Multiplexed coded illumination for Fourier Ptychography with an LED array microscope. Biomedical optics express 5, 2376–2389 (2014).

26 Nguyen, T., Xue, Y., Li, Y., Tian, L. & Nehmetallah, G. Deep learning approach for Fourier ptychography microscopy. Optics express 26, 26470–26484 (2018).

27 Rivenson, Y. et al. Deep learning microscopy. Optica 4, 1437–1443 (2017).

28 Xue, Y., Cheng, S., Li, Y. & Tian, L. Reliable deep-learning-based phase imaging with uncertainty quantification. Optica 6, 618–629 (2019).

29 Nehme, E., Weiss, L. E., Michaeli, T. & Shechtman, Y. Deep-STORM: super-resolution single-molecule microscopy by deep learning. Optica 5, 458–464 (2018).

30 Bayramoglu, N., Kaakinen, M., Eklund, L. & Heikkila, J. in Proceedings of the IEEE International Conference on Computer Vision Workshops. 64–71.

31 Rivenson, Y. et al. PhaseStain: the digital staining of label-free quantitative phase microscopy images using deep learning. Light: Science & Applications 8, 23 (2019).

32 Christiansen, E. M. et al. In silico labeling: predicting fluorescent labels in unlabeled images. Cell 173, 792–803.e719 (2018).

33 Ounkomol, C., Seshamani, S., Maleckar, M. M., Collman, F. & Johnson, G. R. Label-free prediction of three-dimensional fluorescence images from transmitted-light microscopy. Nature methods 15, 917–920 (2018).

34 Fanous, M. et al. Label-free screening of brain tissue myelin content using phase imaging with computational specificity (PICS). Apl Photonics 6, 076103 (2021).

35 Kandel, M. E. et al. Phase Imaging with Computational Specificity (PICS) for measuring dry mass changes in sub-cellular compartments. Nature communications 11, 1–10 (2020).

36 Goswami, N. et al. in Quantitative Phase Imaging VII. 1165313 (International Society for Optics and Photonics).

37 Fanous, M. J., Popescu, G., Tangella, K. & Sobh, N. in Quantitative Phase Imaging VII. 1165311 (International Society for Optics and Photonics).

38 Goswami, N. et al. Label-free SARS-CoV-2 detection and classification using phase imaging with computational specificity. Light: Science & Applications 10, 1–12 (2021).

39 Hu, C. et al. Live-dead assay on unlabeled cells using phase imaging with computational specificity. Nature Communications 13, 713, doi:10.1038/s41467-022-28214-x (2022).

40 Pinkard, H., Phillips, Z., Babakhani, A., Fletcher, D. A. & Waller, L. Deep learning for single-shot autofocus microscopy. Optica 6, 794–797 (2019).

41 Barbastathis, G., Ozcan, A. & Situ, G. On the use of deep learning for computational imaging. Optica 6, 921–943 (2019).

42 de Haan, K. et al. Deep learning-based transformation of H&E stained tissues into special stains. Nature communications 12, 1–13 (2021).

43 Popescu, G. Principles of Biophotonics, Volume 1 - “Linear systems and the Fourier transform in optics”. (IOP Publishing, 2018).

44 Popescu, G. Quantitative phase imaging of cells and tissues. (McGraw Hill Professional, 2011).

45 Isola, P., Zhu, J.-Y., Zhou, T. & Efros, A. A. in Proceedings of the IEEE conference on computer vision and pattern recognition. 1125–1134.

